# Interfering with contextual fear memories by post-reactivation administration of propranolol in mice: a series of null findings

**DOI:** 10.1101/2021.11.01.466717

**Authors:** Wouter R. Cox, Leonidas Faliagkas, Amber Besseling, Rolinka J. van der Loo, Sabine Spijker, Merel Kindt, Priyanka Rao-Ruiz

## Abstract

Post-reactivation amnesia of contextual fear memories by blockade of noradrenergic signaling has been shown to have limited replicability in rodents. This is usually attributed to several boundary conditions that gate the destabilization of memory during its retrieval. How these boundary conditions can be overcome, and what neural mechanisms underlie post-reactivation changes in contextual fear memories remain largely unknown. Here, we report a series of experiments in a contextual fear-conditioning paradigm in mice, that were aimed at solving these issues. We first attempted to obtain a training paradigm that would consistently result in contextual fear memory that could be destabilized upon reactivation, enabling post-retrieval amnesia by administration of propranolol. Unexpectedly, our attempts were unsuccessful to this end. Specifically, over a series of experiments in which we varied different parameters of the fear acquisition procedure, at best small and inconsistent effects were observed. Additionally, we found that propranolol did not alter retrieval-induced neural activity, as measured by the number of c-Fos^+^ cells in the hippocampal dentate gyrus. To determine whether propranolol was perhaps ineffective in interfering with reactivated contextual fear memories, we also included anisomycin (i.e., a potent and well-known amnesic drug) in several experiments, and measures of synaptic glutamate receptor subunit GluA2 (i.e., a marker of memory destabilization). No post-retrieval amnesia by anisomycin and no altered GluA2 expression by reactivation was observed, suggesting that the memories did not undergo destabilization. The null findings are surprising, given that the training paradigms we implemented were previously shown to result in memories that could be modified upon reactivation. Together, our observations illustrate the elusive nature of reactivation-dependent changes of non-human fear memory.

## 1. Introduction

Extensive neuroscientific evidence shows that noradrenergic signaling plays a pivotal role in emotional memory plasticity (Tully and Bolshakov, 2010; McGaugh, 2013; Likhtik and Johansen, 2019). At the occurrence of an emotionally arousing event, noradrenergic transmission in the basolateral amygdala stimulates projections to downstream brain regions, such as the hippocampus, leading to the event being firmly stored in long-term memory (McGaugh, 2015, 2018; Roozendaal and Hermans, 2017). Convincing demonstrations of this crucial involvement of the noradrenergic system in *memory consolidation* come from studies showing that noradrenaline and noradrenergic antagonists, administered shortly before or after fear learning, induce intensified and weakened memory expression, respectively (e.g., Gallagher et al., 1977; Liang et al., 1986; Ji et al., 2003; LaLumiere et al., 2003; Roozendaal et al., 2006; Hu et al., 2007; Johansen et al., 2014; Schiff et al., 2017).

Interestingly, somewhat more recent empirical work has shown that noradrenaline is not only needed for the consolidation of emotional memory, but plays an equally important role in the persistence of memory after its reactivation, a process often referred to as *reconsolidation*. Building on earlier demonstrations of post-reactivation memory loss in the late nineteen-sixties (Misanin et al., 1968), an influential study by Przybyslawski et al. (1999) showed that systemic injections of the β-noradrenergic receptor (β-AR) antagonist propranolol after both encoding and reactivation of emotional memories leads to impaired memory expression 48 h later. Further study of this post-reactivation amnesia seemed to solidify the idea of partly shared underlying mechanisms between memory consolidation and “reconsolidation”. Like the formation of new memories, protein synthesis was shown to be needed for the persistence of memory expression after reactivation (Nader et al., 2000). Hence, it is argued that β-AR antagonists interfere with the creation of new plasticity-related proteins, specifically by preventing noradrenaline from activating the transcription factor cAMP response element-binding protein (CREB) (Thonberg et al., 2002), such that fear memories are not (fully) restabilized in long-term memory (Przybyslawski et al., 1999).

However, the existence of reconsolidation is an issue of considerable debate (Elsey et al., 2018; Gisquet-Verrier and Riccio, 2018). For example, despite the apparent similarities between memory consolidation and reconsolidation, there also seem to be differences in the involved brain regions and circuits (Alberini, 2005; Finnie and Nader, 2012). Furthermore, the mechanisms by which β-AR antagonists interfere with reactivated fear memories remain largely unknown. This is especially true for hippocampus-dependent contextual fear memories, for which the involvement of the noradrenergic system is not as well defined as it is for amygdala-dependent cued fear memories (e.g., Johansen et al., 2014; Schiff et al., 2017). It is evident that synaptic function in the hippocampus can be bidirectionally modulated by the application of adrenergic agonists and antagonists (Gelinas and Nguyen, 2005; Giustino and Maren, 2018). A recent study demonstrated that propranolol has an acute effect on the expression of contextual fear memories and alters memory traces in the dorsal dentate gyrus (DG), basolateral amygdala and pre-frontal cortex when administered before memory reactivation (Leal Santos et al., 2021). However, how blockade of β-ARs may induce amnesia and affect contextual memory traces after drug washout cannot be inferred as no lasting fear-reducing effects of propranolol were found in this study. For these reasons, gaining insights into how exactly β-AR antagonists can impact reactivated contextual fear memories remains of significant interest.

Apart from contributions to the fundamental understanding of memory plasticity, research showing reductions in fear responding by post-reactivation administration of propranolol is relevant in light of clinical applications. A procedure that can be employed to interfere with fear memories would have great utility in the treatment of anxiety disorders and posttraumatic stress disorder. Extinction-based treatments, that currently dominate clinical practice, lead to the creation of a new memory trace that competes with the fear memory. Such procedures do not alter fear memories directly, however, resulting in high relapse rates through a variety of mechanisms (Bouton, 2002). Administering amnestic agents, such as propranolol, in close temporal proximity of memory reactivation seems to result in more durable reductions in fear responding (Dȩbiec et al., 2002; Dȩbiec and Ledoux, 2004; Duvarci and Nader, 2004; Bustos et al., 2006, but see Eisenberg and Dudai, 2004; Lattal and Abel, 2004). Crucially, post-reactivation administration of propranolol has also shown to interfere with fear memories in a human fear-conditioning paradigm (Kindt et al., 2009; Soeter and Kindt, 2010, 2011; Sevenster et al., 2012; Soeter and Kindt, 2012b, 2012a; Sevenster et al., 2013, 2014; Soeter and Kindt, 2015b; Kindt and Soeter, 2018, but see Bos et al., 2014; Schroyens et al., 2017; Chalkia et al., 2019), bringing the development of a new, effective treatment for disorders of emotional memory an important step closer.

Translation of these findings to real-life anxiety disorders and posttraumatic stress disorder remains a great challenge (Kindt, 2014). At present, both promising (Brunet et al., 2008, 2011, 2018; Soeter and Kindt, 2015a; Kindt and van Emmerik, 2016) as well as disappointing (Wood et al., 2015; Elsey et al., 2020; Roullet et al., 2021) results have been reported to treat real-life fears that have been acquired outside the laboratory. These relatively mixed observations have often been attributed to the subtle conditions under which amnestic agents interfere with fear memories (Elsey et al., 2018, 2020). Reactivation of a memory may not be sufficient, but only if a violation of a fear-related expectancy occurs (e.g., not receiving an anticipated aversive stimulus) does administration of amnestic agents lead to reduced fear responding (Pedreira et al., 2004; Lee, 2009; Sevenster et al., 2012). However, with prolonged exposure to the feared stimuli, fear memories could enter a “limbo” state or a new extinction memory is created, such that protein synthesis inhibitors do not target the original fear memory anymore (Merlo et al., 2014; Sevenster et al., 2014). Previous research also suggests that the duration of reactivation at which amnestic agents are effective in reducing subsequent expression of fear intricately depends on memory strength and age, such that older and stronger fear memories may require longer reexposure to the conditioned stimulus (Suzuki et al., 2004; Bustos et al., 2009). This highly complicates translation to clinical practice, as fear memory strength and age vary greatly from patient to patient and currently no measure is available to adjust the reactivation session accordingly. A procedure through which the challenges posed by the boundary conditions could be circumvented would therefore be of great practical value.

Thus, in a quest to (i) elucidate the mechanisms through which β-AR blockers are involved in contextual fear memory reconsolidation and (ii) discover how boundary conditions of post-reactivation amnesia can be overcome, we performed a series of experiments in an animal fear-conditioning paradigm. Towards these goals, our first aim was to obtain a behavioral paradigm that would consistently show reactivation-induced amnesia by the administration of a β-AR antagonist after contextual fear memory reactivation. We performed a series of experiments (including replications) with varied training protocols, and post-reactivation administration of propranolol. However, these attempts to induce robust post-reactivation amnesia were unsuccessful, such that no further experimentation was possible. Our behavioral data sharply contradict earlier successful reports (Abrari et al., 2008; Muravieva and Alberini, 2010; Liu et al., 2015), but are reminiscent of a recent series of likewise unsuccessful experiments in rats that includes the administration of different amnesic agents (i.e., propranolol, midazolam, and cycloheximide) (Schroyens et al., 2019). In line with these data, we demonstrate – using a different species – that post-reactivation amnesia of contextual fear memories by systemic administration of propranolol does not work with flawless fidelity on both a behavioral (freezing) and cellular level (reactivation-induced activity of putative memory trace cells in the hippocampal dentate gyrus). In addition, no effect of the commonly used amnestic agent anisomycin (Barbacid and Vazquez, 1974), and no reactivation-dependent regulation of molecular markers (downregulation of the synaptic glutamate receptor subunit GluA2) was found, indicating that we were unable to trigger memory destabilization and subsequent restabilization (Rao-Ruiz et al., 2011; Bhattacharya et al., 2017) using training protocols that were previously successful in this respect (Rao-Ruiz et al., 2011). Together, the behavioral, cellular, and molecular data provided here comprises a valuable resource for future research aimed at elucidating the precise conditions under which contextual fear memories can be destabilized and manipulated upon their reactivation.

## 2. Materials and Methods

### 2.1. General procedures

#### 2.1.1. Animals

Male C57BI6/J inbred mice (8–10 weeks of age), from Charles River France, were individually housed with enrichment and kept on a 12/12 h light/dark cycle (7 A.M. lights on) with food and water available ad libitum. The mice acclimatized to their home cage for two weeks prior to experimentation. All procedures were carried out during the light phase, between 9 A.M. and 12 P.M., unless otherwise specified. Prior to the start of the experiment, mice underwent three consecutive days of handling (2–3 minutes per mouse) to habituate the animals to being held and restrained by the experimenter. All experiments were performed in accordance with Dutch law and licensing agreements using protocols ethically approved by the Animal Ethical Committee of the VU University Amsterdam.

#### 2.1.2. Apparatus

The experiments were executed in one of two automated fear-conditioning systems: (i) TSE Fear Conditioning System for Small Rodents (TSE systems, Germany) in Experiments 1–2 (e.g., Rao-Ruiz et al., 2011; Végh et al., 2014) and (ii) Med Associates Video Fear Conditioning System (Sandown Scientific, UK) in Experiments 3–9 (e.g., Anagnostaras et al., 2010; Gouty-Colomer et al., 2016). The mice underwent contextual fear conditioning, memory reactivation, and a retention test in a conditioning chamber with a stainless-steel grid floor, constant house light (TSE: 100–500 lx, Med Associates: White light: 450–650 nm and near-infrared light: 940 nm) and background white noise (TSE: 68 dB, Med Associates: 50 dB, 5000 Hz). The fear-conditioning box was placed in a sound attenuating outer chamber. The apparatus was cleaned with 70% ethanol prior to each session.

#### 2.1.3. Drugs

Propranolol HCl (Sigma-Aldrich) was dissolved in saline at a ratio of 2 mg/mL, unless otherwise specified, and anisomycin (Sigma-Aldrich) at a ratio of 30 mg/1500 mL. The mice in the experimental groups received 10 mg per kg body weight of propranolol (Przybyslawski et al., 1999; Villain et al., 2016) or anisomycin at 150 mg per kg body weight (Suzuki et al., 2004; Rao-Ruiz et al., 2011). The mice in the control group received the same volume of saline. The drugs were administered immediately after memory reactivation through intraperitoneal injections in all experiments.

#### 2.1.4. Data analysis

Freezing, defined as the cessation of all movement except respiration and heartbeat, was used as a measure of fear memory expression, and scored using automated algorithms in the TSE (Rao-Ruiz et al., 2011) or Med Associates (Anagnostaras et al., 2010; Gouty-Colomer et al., 2016) systems. For the TSE chamber (Experiments 1–2), a time threshold of 1 s was used for automatically recorded “freezing” behavior (i.e., no detected change in sensor status for 1 s in X, Y and Z axes). For the Med associates chamber (Experiments 3–9), freezing was measured by Video Freeze Software (Anagnostaras et al., 2010) using a linear method with a motion threshold of 18 a.u., sample rate of 30 frames per second and a minimum freeze duration of 30 frames. Freezing data is presented as a percentage of total session time.

Statistical analyses were carried out using the Statistical Package for the Social Sciences (SPSS) version 25.0 (Armonk, NY: IBM Corp.). To assess whether the administration of amnestic drugs (propranolol or anisomycin) after memory reactivation results in an attenuation of fear responding, we performed a mixed ANOVA with Drug (amnestic drug *vs*. saline) as between-subjects factor and Session (reactivation, retention) as within-subjects factor. When relevant, we performed additional independent t-tests. To make sure that the null findings cannot be attributed to any animals not having acquired contextual fear, we performed additional analyses in which only animals exhibiting a freezing level of 10% or higher during reactivation were included (see Figure S1). This inclusion of mice based on freezing levels did not lead to different results. Exclusion of outliers or non-parametric analyses of experiments in which assumptions of normality or homogeneity of variances were violated did not change any results either.

### 2.2. Specific procedures

#### 2.2.1. Experiment 1

##### 2.2.1.1. Contextual fear conditioning

Mice were placed in the fear-conditioning chamber, and after 180 s, received a 2 s 0.7 mA foot shock as the unconditioned stimulus (US). Thirty seconds later, the mice were removed from the box and placed in their home cage.

##### 2.2.1.2. Memory reactivation

Twenty-four hours afterwards, to reactivate the fear memory, the mice were placed in the conditioning box for 180 s without delivery of the US. Immediately thereafter, the mice were weighed and received either saline or propranolol in a quasi-random fashion (i.e., alternation) and then returned to their home cage.

##### 2.2.1.3. Retention test

Forty-eight hours after memory reactivation (Dȩbiec & Ledoux, 2004), mice were placed in the conditioning box for 180 s in the absence of the unconditioned stimulus and returned to their home cage immediately afterwards. One mouse was excluded due to a procedural error (Experiment 1b, saline condition).

##### 2.2.1.4. Immunohistochemistry

Ninety minutes after the retention test, 5 animals from each experimental group (saline and propranolol), were randomly selected and transcardially perfused with ice-cold PBS, followed by 4% paraformaldehyde (Sigma-Aldrich Chemie N.V., The Netherlands). Forty micrometer coronal sections of the hippocampus were collected serially using a freezing microtome (Leica, Wetzlar, Germany; SM 2000R) and stored in PBS with 0.02% NaN3 at 4 °C until further use. Approximately 8–10 free floating sections across the rostrocaudal axis of the dorsal DG were used for immunohistochemical stainings (described in Rao-Ruiz et al., 2019) using a primary antibody against the Immediate Early Gene c-Fos (1:500, sc-52, Santa Cruz, Germany) and a corresponding Alexa-conjugated secondary antibody (1:400, anti-rabbit Alexa 488, Life Technologies, The Netherlands). Nuclear staining was performed using DAPI (300 nmol/L, Thermo Fisher Scientific, The Netherlands). Sections were mounted on slides and coverslipped using Polyvinyl alcohol mounting medium with DABCO®, antifading (Merck KGaA, Germany).

##### 2.2.1.5. Confocal microscopy, cell counting and data analysis

A Nikon Instruments A1 Confocal Laser Microscope with NIS-Elements C Software was used to make approximately 8–10 z-stacks of the DG/animal at 10x magnification by an experimenter blinded to the experimental condition. Images were imported to Fiji (Version 1.0) where they were digitally merged to form composite images. Individual c-Fos^+^ cells in the DG were manually marked, counted using the Cell Counter plugin in Fiji, and averaged/1.3 mm section for each animal. Representative images were edited in ImageJ to generate 2D projections of z-stacks, and all images were treated identically.

#### 2.2.2. Experiment 2

##### 2.2.2.1. Contextual fear conditioning

During fear conditioning, a 1 mA foot shock was administered (instead of a 0.7 mA foot shock as described in Experiment 1). All other procedures were the same as in Experiment 1.

##### 2.2.2.2. Immunohistochemistry

Seven animals randomly selected from each experimental group (saline and propranolol) were used for c-Fos immunohistochemistry as described in Experiment 1.

#### 2.2.3. Experiment 3

In addition to administering propranolol in the same way as in Experiment 1– 2, we included a condition in which this drug was dissolved at a ratio of 1 mg/1 mL (Villain et al., 2016), and injected with double the volume. All other procedures were the same as in Experiment 2.

#### 2.2.4. Experiment 4

The mice were kept on a reversed, 12 h light/dark scale, such that all experimental procedures (i.e., habituation, handling, conditioning, reactivation, retention testing) took place during the dark phase, between 9 A.M. and 12 P.M. All other behavioral procedures were the same as in Experiment 2.

#### 2.2.5. Experiment 5

##### 2.2.5.1. Background contextual fear conditioning

The mice were placed in the conditioning chamber, and after 120 s, a tone (5 kHz, 75 dB) was presented for 30 s, followed by a 2 s 1 mA foot shock. Thirty seconds after shock discontinuation, the mice were removed from the conditioning chamber and returned to their home cage.

##### 2.2.5.2. Memory reactivation

Forty-eight hours after conditioning, mice were placed back in the conditioning box for 90 s without tone or shock delivery.

##### 2.2.5.3. Retention test

Forty-eight hours after reactivation, mice were placed in the conditioning box for 180 s without delivery of a tone or shock, as a test of contextual fear.

#### 2.2.6. Experiment 6

In addition to propranolol, we included a condition in which animals received anisomycin. All other behavioral procedures were the same as in Experiment 2.

#### 2.2.7. Experiment 7

##### 2.2.7.1. Contextual fear conditioning

During fear conditioning, no shock, one 0.7 mA shock, or one 1 mA foot shock was administered. All other procedures were the same as in Experiment 6.

##### 2.2.7.2. Memory reactivation

Mice were sacrificed via cervical dislocation (Rao-Ruiz et al., 2011) 1 hour after memory reactivation, and propranolol or saline administration,. All other procedures were the same as in Experiment 6.

##### 2.2.7.3. Brain tissue preparation

Dorsal hippocampi were dissected on ice, snap frozen and stored at -80 °C until synaptosome isolation.

##### 2.2.7.4. Synaptosome isolation

Synaptosomes were isolated on a sucrose gradient as described previously (Pandya et al., 2017). Briefly, hippocampi from each animal were homogenized in ice-cold Homogenization buffer (2 M sucrose, 500 mM HEPES (pH 7,4)) containing complete Protease Inhibitor (Roche/Sigma) and spun down for 10 min at 1000 x g at 4 °C. The supernatant was layered on top of a 0.85 M/1.2 M sucrose gradient, spun at 100,000 x g for 2h at 4 °C and synaptosomes were recovered at the interface of 0.85/1.2 M sucrose. Protein concentration was determined using a Bradford assay (BioRad).

##### 2.2.7.5. Immunoblot analysis

Immunoblot analysis was performed as described previously (Gonzalez-Lozano et al., 2021) on 5 µg of synaptosomal protein from each experimental group. After electrophoresis, gels were scanned using a Gel Doc EZ imager (Bio-Rad) and electro-transferred onto a PVDF membrane overnight at 40 V. After blocking in 5% milk, membranes were incubated with a primary antibody against GluA2 (1:0000, Cat nr. 75-002, Neuromab) overnight at 4 °C and with a matching HRP-conjugated secondary antibody for 2 h at room temperature. The membranes were then scanned with Femto ECL Substrate (Thermo Fisher Scientific, Waltham, MA, USA) on an Odyssey Fc system (LI-COR Bioscience, Lincoln, NA, USA) and quantified using Image Studio software (version 2.0.38). Input differences were corrected using total protein amounts loaded on the gel (Gonzalez-Lozano et al., 2021).

#### 2.2.8. Experiment 8

##### 2.2.8.1. Contextual fear conditioning

The procedure was based on an earlier study that has shown post-reactivation memory loss in rats (Schmidt et al., 2017). During conditioning, the mice were placed in the conditioning chamber and, after 120 s, received a total of three 0.5 mA shocks with a 2 s duration. The shocks were administered with regular intervals of 30 s in between each shock. Thirty seconds after discontinuation of the last US, the mice were removed from the conditioning chamber, and returned to their home cage. All other procedures were the same as in Experiment 6.

#### 2.2.9. Experiment 9

##### 2.2.9.1. Contextual fear conditioning

Instead of three 0.5 mA shocks, the mice received three 0.3 mA shocks. All other procedures were the same as in Experiment 8.

## 3. Results

### 3.1. Experiment 1

In Experiment 1, we tested whether the administration of propranolol after reactivation of a contextual fear memory eliminates expression of fear during a retention test 48 h later, using a behavioral protocol that has been shown to result in post-reactivation amnesia by anisomycin (Suzuki et al., 2004; Rao-Ruiz et al., 2011) (Figure 1A). We observed that propranolol did not lead to reduced fear responding (Drug × Session, *F*1,18 = 0.021, *p* = 0.887, Figure 1B). This finding is not in line with post-reactivation amnesia using propranolol in previous studies (e.g., Taherian et al., 2014). Therefore, to ascertain that the present finding was not a false negative, we performed an exact replication of experiment 1a (Experiment 1b).

**Figure. 1.**
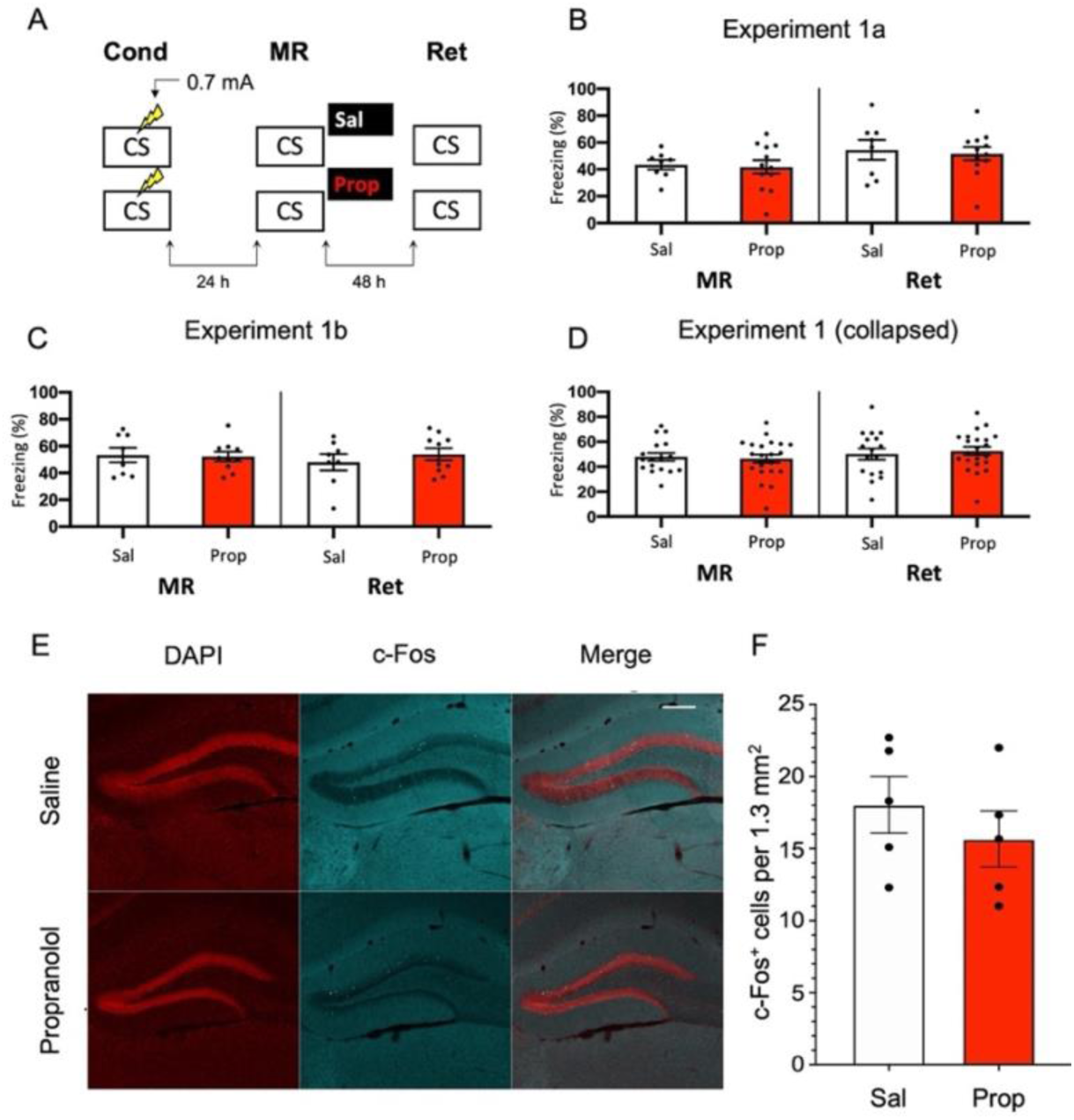
No effect of propranolol on reactivated memories that were acquired with a single 0.7 mA shock during contextual fear conditioning. **(A)** Design of Experiment 1. Mice received one 0.7 mA shock during fear conditioning. One day later, saline (n = 8) or propranolol (n = 12) was administered after memory reactivation. Forty-eight hours afterwards, retention of contextual fear was assessed. Cond = conditioning, MR = memory reactivation, Ret = retention, CS = conditioned stimulus (context), Sal = saline, Prop = propranolol. **(B)** Results of Experiment 1a. Average percentage of freezing during memory reactivation is displayed on the left, and freezing during the retention test on the right panel of the column chart (saline in white bars, propranolol in red bars). Error bars represent SEM. Filled circles indicate individual animals. **(C)** Results of Experiment 1b (saline, n = 9; propranolol, n = 10). Experiment 1b was a direct replication of Experiment 1a. **(D)** Collapsed results of Experiment 1a and Experiment 1b. **(E)** Representative images of the DG 90 minutes after the retention test from mice that received either saline (n =5), or propranolol (n=5) after reactivation (Experiment 1a). Cells that were c-Fos+ are labelled in cyan (scale bar: 200 μm). **(F)** Number of c-Fos+ cells in the DG per 1.3 mm2. Error bars represent SEM. Filled circles indicate individual animals.

Like Experiment 1a, in Experiment 1b there was no effect of propranolol on the pattern of freezing levels from memory reactivation to the retention test two days later (Drug × Session, *F*1,16 = 0.582, *p* = 0.457, Figure 1C). To ensure that the two present null findings are not a reflection of insufficient power, we collapsed the data of Experiment 1a and 1b, and repeated the analysis. No effect was observed on the collapsed freezing data either (*F*1,36 = 0.277, *p* = 0.602, Figure 1D).

We also tested if noradrenergic signaling after memory reactivation affected retrieval-induced neural activity in the DG, where memory traces for contextual fear have been previously identified (Liu et al., 2012; Denny et al., 2014; Rao-Ruiz et al., 2019). To this end, we measured the number of c-Fos^+^ cells, a molecular correlate of neural activity, 90 minutes after the retention test in the granule cell layer of the DG. In line with our behavioral results, no significant differences were observed in the number of activated DG neurons (c-Fos^+^) between experimental groups (*t*8 = 0.856, *p* = 0.417, Figures 1 E and F). Together, these data indicate that post-reactivation administration of propranolol does not affect subsequent memory retention or the activation of hippocampal neurons that are considered to be the cellular substrate of memory using the current experimental protocol.

### 3.2. Experiment 2

Since previous research has shown (though not consistently, Taherian et al., 2014) that memory strength modulates post-reactivation amnesia (Suzuki et al., 2004), we hypothesized that fear acquisition in Experiment 1 may not have been robust enough with the current protocol. Indeed, close inspection of the data of Experiment 1 shows that the freezing levels were considerably lower than in a previous study from our lab, which involved the same mouse strain, laboratory space, behavioral system and training and memory reactivation protocols (Rao-Ruiz et al., 2011, for accurate comparison see Figure S2 for the results of Experiment 1 using the same time threshold for freezing behavior as in this previous study). To induce stronger fear memories, we administered a single, higher intensity, i.e. 1 mA, foot shock during conditioning (Figure 2A).

**Figure. 2.**
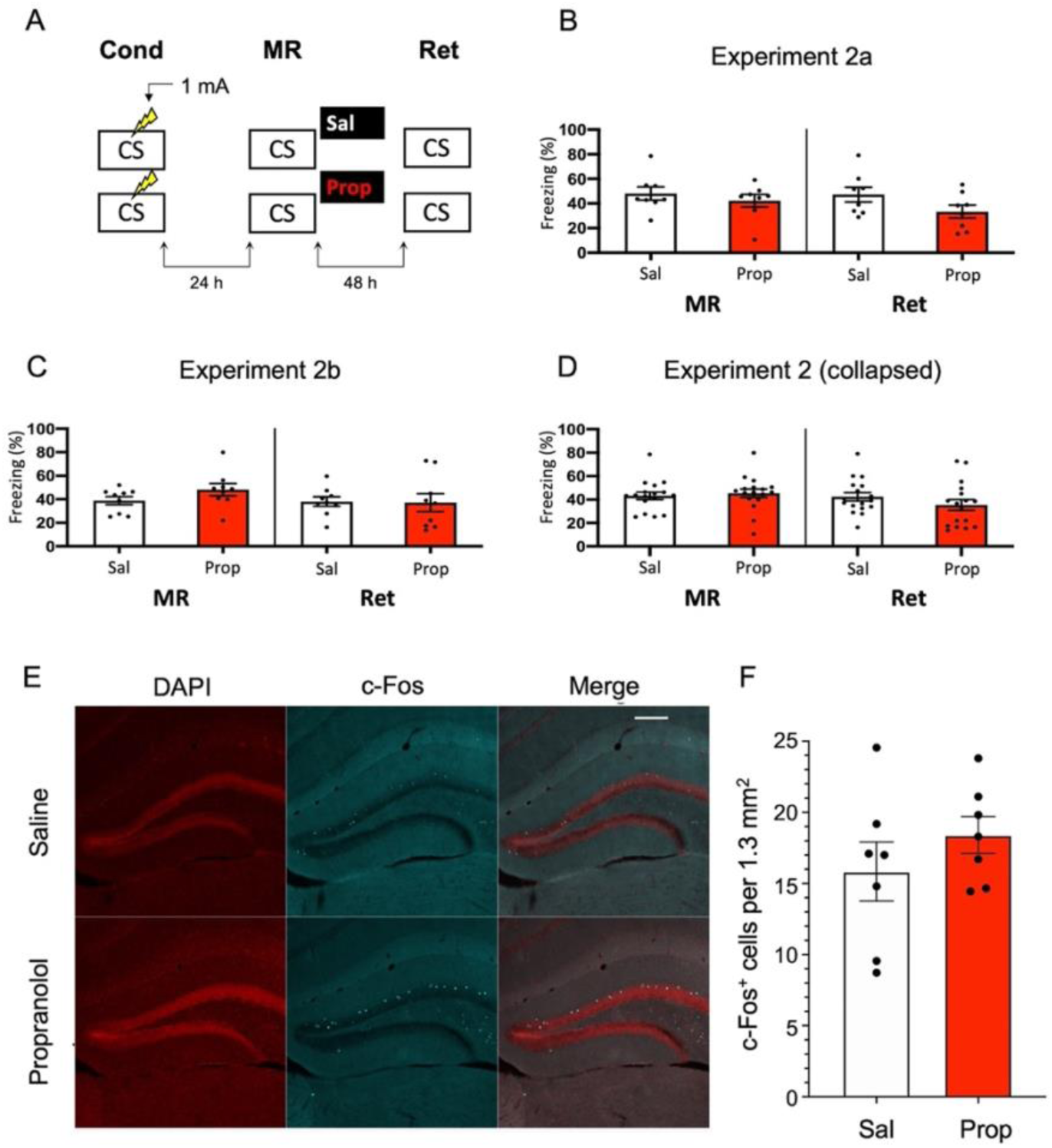
A minor effect of propranolol on reactivated memories that were acquired with a single 1 mA shock during contextual fear conditioning. (**A**) Design of Experiment 2a. Mice received one 1 mA shock during fear conditioning. One day later, saline (*n* = 8) or propranolol (*n* = 8) was administered after memory reactivation. Forty-eight hours afterwards, retention of contextual fear was assessed. Cond = conditioning, MR = memory reactivation, Ret = retention, CS = conditioned stimulus (context), Sal = saline, Prop = propranolol. (**B**) Results of Experiment 2a. Average percentage of freezing during memory reactivation is displayed on the left, and freezing during the retention test on the right panel of the column chart (saline in white bars, propranolol in red bars). Error bars represent SEM. Filled circles indicate individual animals. (**C**) Results of Experiment 2b (saline, *n* = 9; propranolol, *n* = 9). Experiment 2b was a direct replication of Experiment 2a. (**D**) Collapsed results of Experiment 2a and Experiment 2b. (**E)** Representative images of the DG 90 minutes after the retention test from mice that received either saline (n =7), or propranolol (n=7) post-reactivation (Experiment 2a). Cells that were c-Fos^+^ are labelled in cyan (scale bar: 200 μm). (**F**) Number of c-Fos^+^ cells in the DG per 1.3 mm^2^. Error bars represent SEM. Filled circles indicate individual animals.

However, no reduction in fear responding by propranolol was found in this experiment either (Drug × Session, *F*1,14 = 1.200, *p* = 0.292, Figure 2B). To make sure that the results were not a false negative, we repeated the experiment (Experiment 2b). Again, no significant effect was observed (Drug × Session, *F*1,16 = 2.523, *p* = 0.132, Figure 2C). When the data of Experiment 2a and 2b were collapsed we did find a trend for reduction in freezing from memory reactivation to retention test in the propranolol relative to the saline condition (Drug × Session, *F*1,32 = 3.827, *p* = 0.059, Figure 2D). Notably, raising the foot shock level in Experiment 2 did not lead to higher freezing levels during reactivation than in Experiment 1 (*t*70 = 0.876, *p* = 0.384, compare Figures 1D and 2D). It also bears mentioning that this reduction in fear responding by post-reactivation administration of propranolol after collapsing the data of Experiment 2a and 2b is considerably smaller than in previous studies (e.g., Liu et al., 2015; Villain et al., 2016).

Nevertheless, since a marginal amnestic effect of propranolol was observed, we further tested if neural activity in the DG was affected as well. However, as in Experiment 1, administration of propranolol had no effect on the number of DG neurons activated (c-Fos^+^) by the retention test (*t*12 = 1.046, *p* = 0.316, Figure 2E and F). We reasoned that the lack of a robust effect of post-reactivation propranolol on the retention test could be due to two factors: (i) limited efficacy of propranolol, or (ii) relatively unsuccessful triggering of memory destabilization. These two explanations were addressed in the subsequent experiments (3–5, and 6, respectively).

### 3.3. Experiment 3

As propranolol has lipophilic properties (Cruickshank, 1980), it is possible that a substantial amount of the drug binds to body fat upon injecton, such that noradrenergic signaling in the brain was only partially blocked in Experiment 2, resulting in reduced efficacy and relatively mild amnesia. Therefore, we performed an experiment that was the same as Experiment 2, but now also included a condition in which propranolol was dissolved in saline at half the ratio and injected in double the volume/body weight (i.e., to reduce binding to body fat without changing the dose). However, injecting propranolol in a larger volume of saline did not lead to a more robust decrease in fear responding (Drug × Session, *F*2,24 = 0.499, *p* = 0.613, Figure 3A, B).

**Figure. 3.**
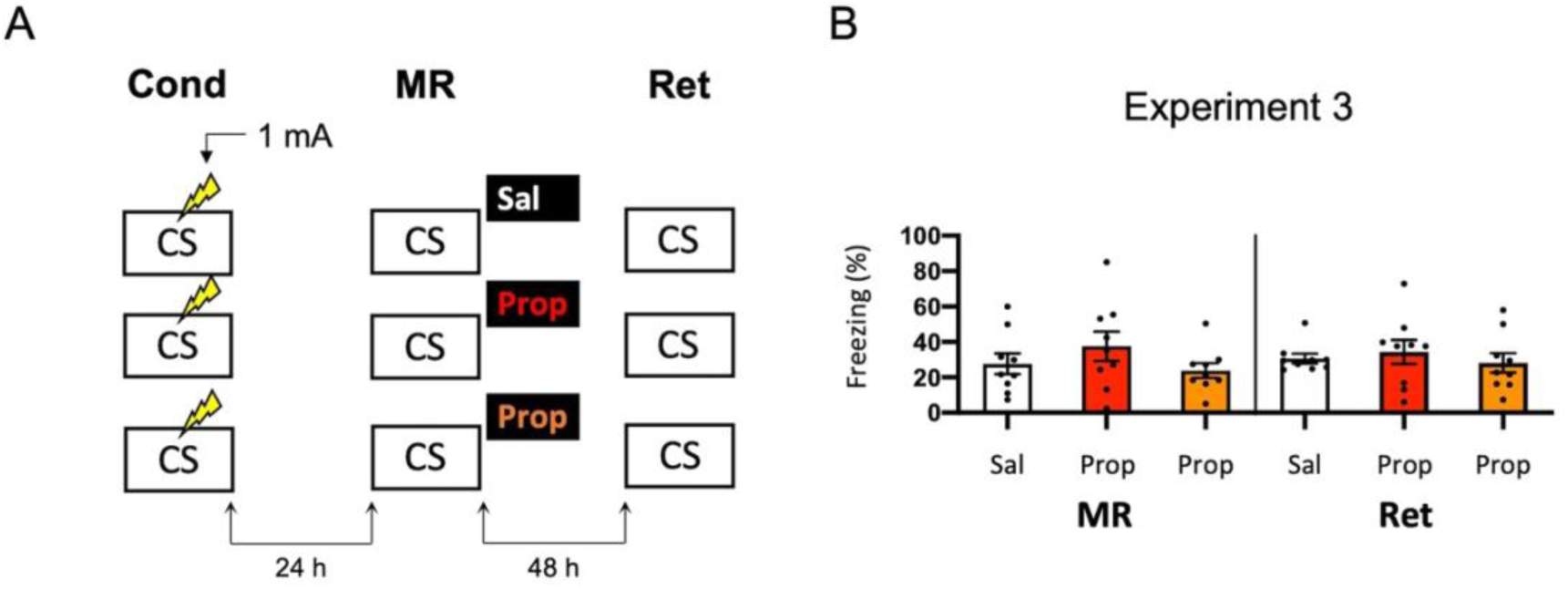
No effect of propranolol – injected in two different volumes – on reactivated memories that were acquired with a single 1 mA shock during contextual fear conditioning. (**A**) Design of Experiment 3. Mice received one 1 mA shock during fear conditioning. One day later, mice received saline (*n* = 9), propranolol (*n* = 9), or propranolol dissolved in saline at half the ratio and injected in double the volume/body weight (*n* = 9) after memory reactivation. Forty-eight hours afterwards, retention of contextual fear was assessed. Cond = conditioning, MR = memory reactivation, Ret = retention, CS = conditioned stimulus (context), Sal = saline, Prop = propranolol. (**B**) Results of Experiment 3. Average percentage of freezing during memory reactivation is displayed on the left, and freezing during the retention test on the right panel of the column chart (saline in white bars, propranolol in red bars, and propranolol double volume in orange bars). Error bars represent SEM. Filled circles indicate individual animals.

### 3.4. Experiment 4

Another potential explanation for a small effect of propranolol in Experiment 2 might be that the experiments were performed during the light phase (i.e., the sleep cycle of mice). Previous research has shown that noradrenergic signaling from the locus coeruleus goes into time-outs during specific phases of the sleep cycle (Aston-Jones and Bloom, 1981), such that synaptic plasticity is impaired (Cirelli and Tononi, 2000). Therefore, in Experiment 2 the memories might have been reactivated and restabilized with little involvement of those receptors that propranolol specifically targets (i.e., β-adrenergic receptors, Johansen et al., 2011). Hypothetically, a more robust effect of propranolol on reactivated fear might be observed when memories are acquired and reactivating in the animals’ wake-phase. Since rodents are nocturnal, we repeated Experiment 2, but now performed all experimental procedures in the animals’ dark phase (Figure 4A). Again, no fear reduction by propranolol was found (Drug × Session, *F*1,22 = 0.031, *p* = 0.861, Figure 4B). The freezing levels during memory reactivation were notably low compared to the previously performed experiments (e.g., Experiment 2), even after exclusion of mice with freezing levels below 10% (Figure S1H). This may be attributable to light enhancing freezing levels (Warthen et al., 2011), so that when experiments are performed in the dark phase the animals freeze to a particularly low degree.

**Figure. 4.**
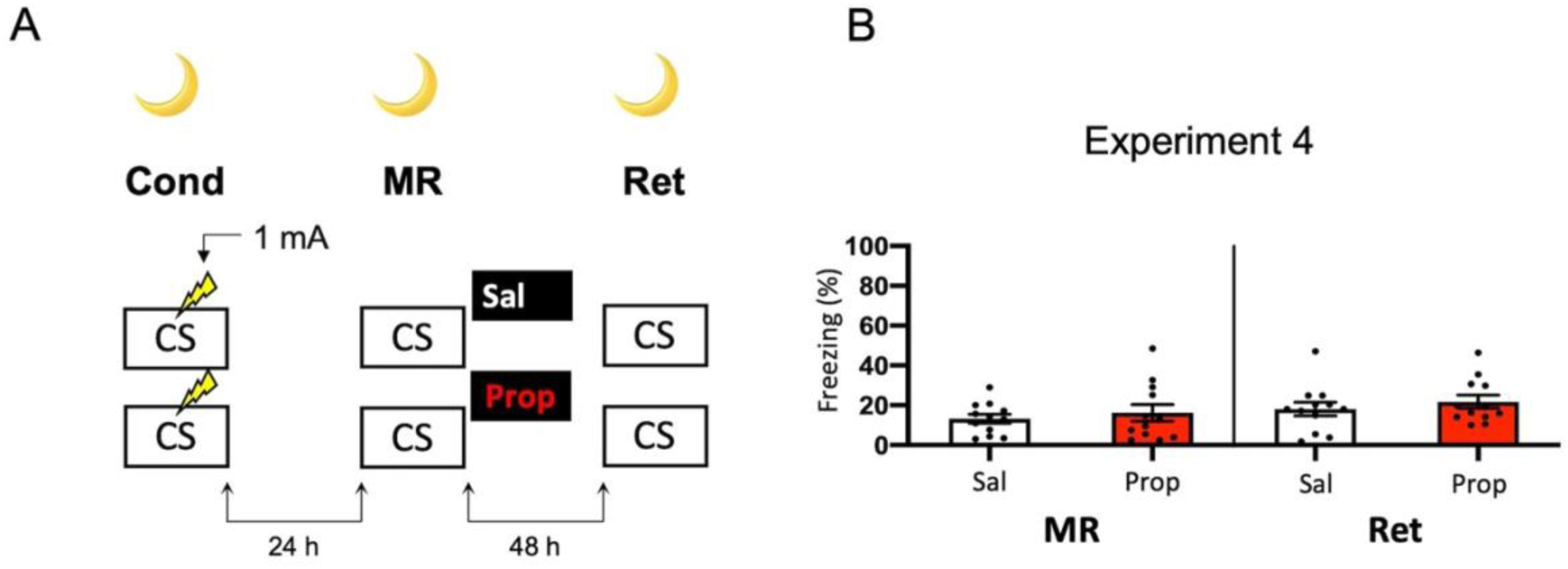
No effect of propranolol on reactivated memories that were acquired with a single 1 mA shock during contextual fear conditioning in the animals’ dark phase. (**A**) Design of Experiment 4. Mice received one 1 mA shock during fear conditioning. One day later, mice received saline (*n* = 12) or propranolol (*n* = 12) after memory reactivation. Forty-eight hours afterwards, retention of contextual fear was assessed. All procedures took place in the animals’ dark phase. Cond = conditioning, MR = memory reactivation, Ret = retention, CS = conditioned stimulus (context), Sal = saline, Prop = propranolol. (**B**) Results of Experiment 4. Average percentage of freezing during memory reactivation is displayed on the left, and freezing during the retention test on the right panel of the column chart (saline in white bars, propranolol in red bars). Error bars represent SEM. Filled circles indicate individual animals.

### 3.5. Experiment 5

A last potential way to increase the efficacy of post-reactivation administration propranolol and to induce more robust amnesia could be to alter the way in which the contextual fear memory is processed. For example, previous research has shown differences in how foreground contextual fear conditioning and background contextual fear conditioning are encoded and the role of the hippocampus in each (Trifilieff et al., 2006). Furthermore, a study has shown post-reactivation amnesia with propranolol in a background contextual fear-conditioning paradigm (Muravieva and Alberini, 2010). In the next experiment, we therefore aimed to induce amnesia by administering propranolol after reactivation of a memory that was acquired by background contextual fear conditioning (Figure 5A). However, the results showed no effect of propranolol on contextual fear (Drug × Session, *F*1,20 = 1.591, *p* = 0.222, Figure 5B).

**Figure. 5.**
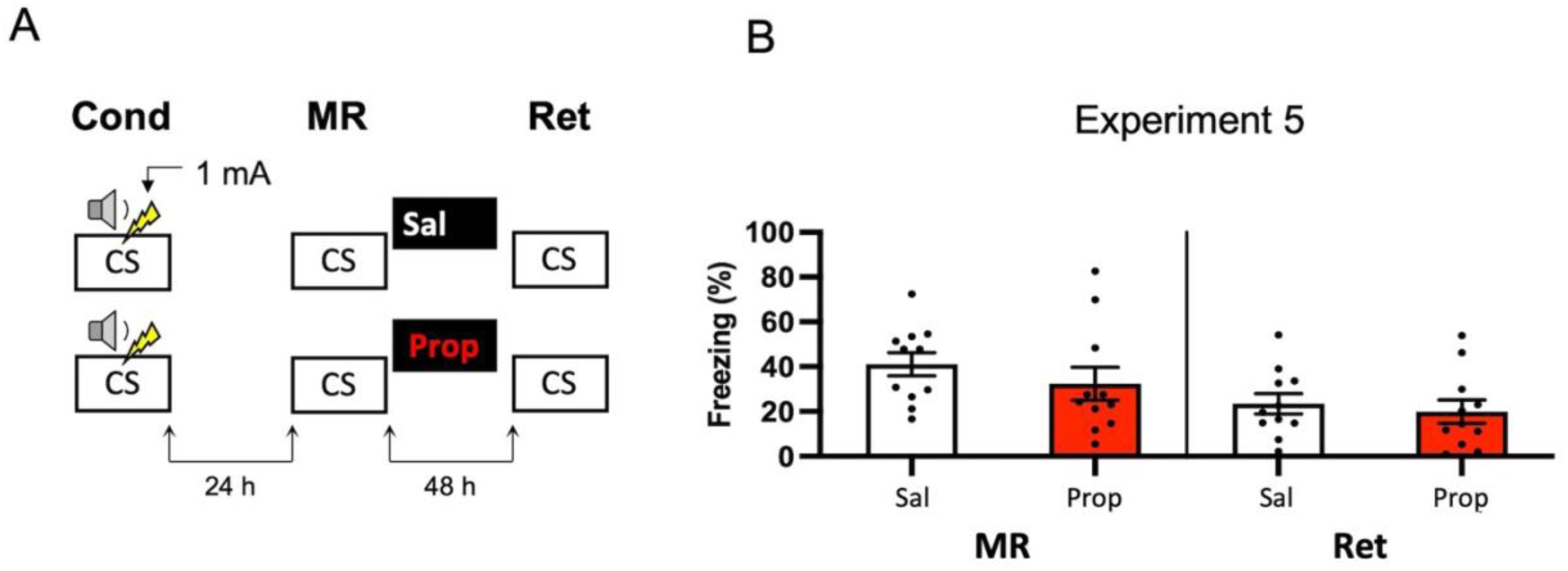
No effect of propranolol on reactivated memories in a background contextual fear-conditioning paradigm. (**A**) Design of Experiment 5. Mice received one 1 mA shock during presentation of a tone. One day later, mice received saline (*n* = 11) or propranolol (*n* = 11) after contextual memory reactivation. Forty-eight hours afterwards, retention of contextual fear was assessed by exposure to the context. Cond = conditioning, MR = memory reactivation, Ret = retention, CS = conditioned stimulus (context), Sal = saline, Prop = propranolol. (**B**) Results of Experiment 5. Average percentage of freezing during memory reactivation is displayed on the left panel of the column chart, and freezing during the retention test on right panel of the column chart (saline in white bars, propranolol in red bars). Error bars represent SEM. Filled circles indicate individual animals.

### Experiment 6

In Experiments 3–5, we aimed to obtain a more pronounced reduction in fear responding by post-reactivation administration of propranolol than in Experiment 2 by studying potential ways that could increase the efficacy of propranolol. We did so by altering the volume in which propranolol is injected (Experiment 3) and creating contextual fear memories that could be more sensitive to noradrenergic signaling (Experiments 4–5). Since no amnestic effects were observed, we reasoned that the null findings may not be caused by an inefficacy of propranolol, but rather unsuccessful induction of memory destabilization. To shed light on this idea, we repeated Experiment 2 but now included a condition in which anisomycin, a potent inhibitor of eukaryotic protein synthesis (Barbacid and Vazquez, 1974) and one of the most widely studied amnestic agents in the reconsolidation literature (e.g., Nader et al., 2000; Lee et al., 2004; Suzuki et al., 2004; Parsons et al., 2006; Blundell et al., 2008; Rao-Ruiz et al., 2011; Kwak et al., 2012) was administered (Figure 6A). No reduction in fear responding by either anisomycin or propranolol was found (Drug × Session, *F*2,22 = 0.327, *p* = 0.724, Figure 6B). Furthermore, a follow-up analysis with the collapsed data of the saline and propranolol conditions in Experiment 3 and 6 showed no significant amnesic effects of propranolol (*F*1,42 = 0.530, *p* = 0.471), unlike the pooled data of Experiment 2 in which the same procedure was followed (compare Figures 3B and 6B *vs*. Figure 2D). These findings suggest that the lack of post-reactivation amnesia that we observe seems attributable to an ineffectiveness of the memory reactivation session in triggering memory destabilization and hence its subsequent reconsolidation process.

**Figure. 6.**
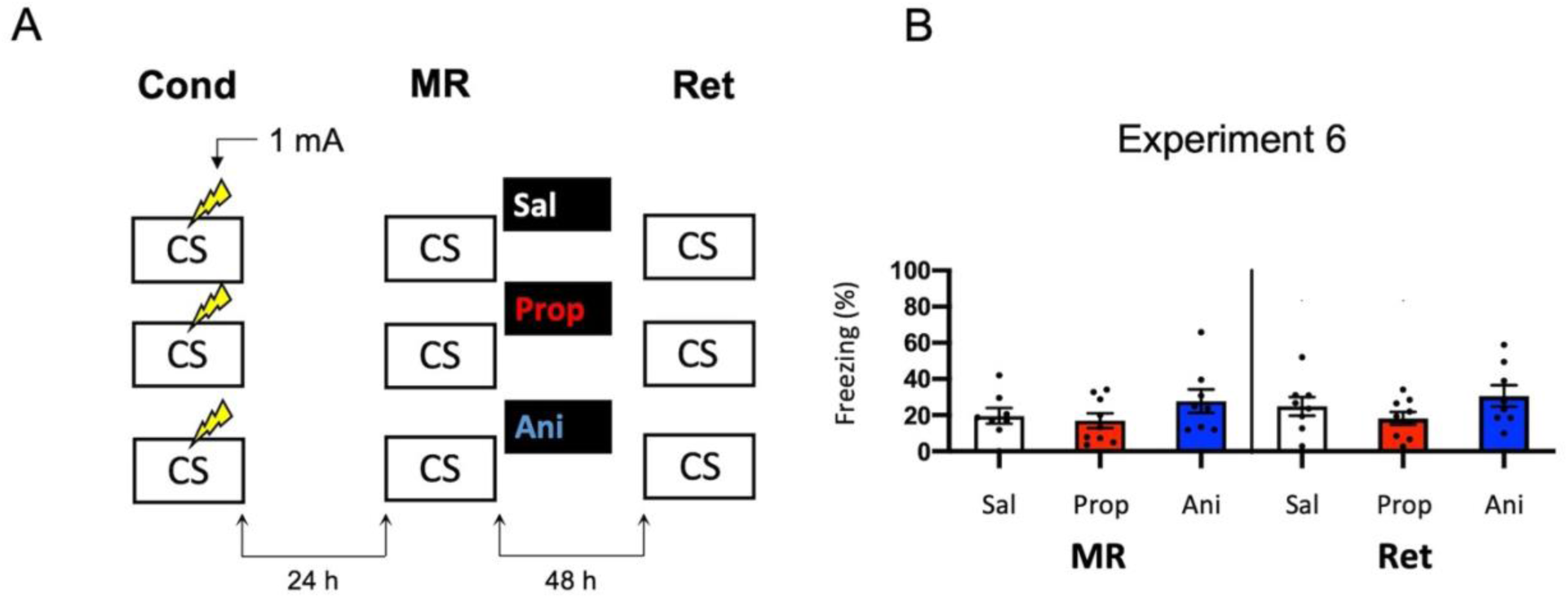
No effect of propranolol or anisomycin on reactivated memories that were acquired with a single 1 mA shock during contextual fear conditioning. (**A**) Design of Experiment 6. Mice received one 1 mA shock during fear conditioning. One day later, mice received saline (*n* = 8), propranolol (*n* = 9), or anisomycin (*n* = 8) after memory reactivation. Forty-eight hours afterwards, retention of contextual fear was assessed. Cond = conditioning, MR = memory reactivation, Ret = retention, CS = conditioned stimulus (context), Sal = saline, Prop = propranolol, Ani = anisomycin. (**B**) Results of Experiment 6. Average percentage of freezing during memory reactivation is displayed on the left, and freezing during the retention test on the right panel of the column chart (saline in white bars, propranolol in red bars, and anisomycin in blue bars). Error bars represent SEM. Filled circles indicate individual animals.

### 3.6. Experiment 7

We - and others - have previously shown that synaptic trafficking of the glutamate receptor subunit GluA2 in the hippocampus is crucially involved in contextual fear memory destabilization (Rao-Ruiz et al., 2011; Bhattacharya et al., 2017). Specifically, upon memory reactivation, endocytosis of GluA2-containing receptors – leading to downregulation of synaptic GluA2 subunits – is needed for retrieval-induced memory plasticity. Therefore, reduced expression of GluA2 shortly after memory reactivation can be used as a marker of destabilization (Rao-Ruiz et al., 2011, Bhattacharya et al., 2017). In Experiment 7, we made use of this molecular marker to gain additional insight into whether, in the present series of experiments, propranolol has been ineffective in interfering with reactivated memories or rather that the protocols have not induced memory destabilization. Mice underwent contextual fear conditioning including a single 0.7 mA shock or a 1 mA shock and were exposed to the conditioning context 24 hours later using the same protocols as before (Experiment 1–3, 6) (Figure 7A). An additional group of mice, that served as baseline controls, were exposed to the conditioning context but did not receive shocks during the conditioning phase (Figure 7A). One hour after memory reactivation and treatment with propranolol or saline, synaptic GluA2 expression was assessed using immunoblot analysis (Rao-Ruiz et al., 2011). Neither the 0.7 mA, nor the 1 mA shock groups showed a significant downregulation of GluA2 subunits in dorsal hippocampal synapses, relative to no shock controls (Shock, *F*2,27 = 0.130, *p* = 0.879, Figure 7B, C). This finding thus indicates that our present null findings are not due to ineffectiveness of propranolol as an amnestic drug, but that the reactivation sessions did not trigger memory destabilization and subsequent reconsolidation.

**Figure. 7.**
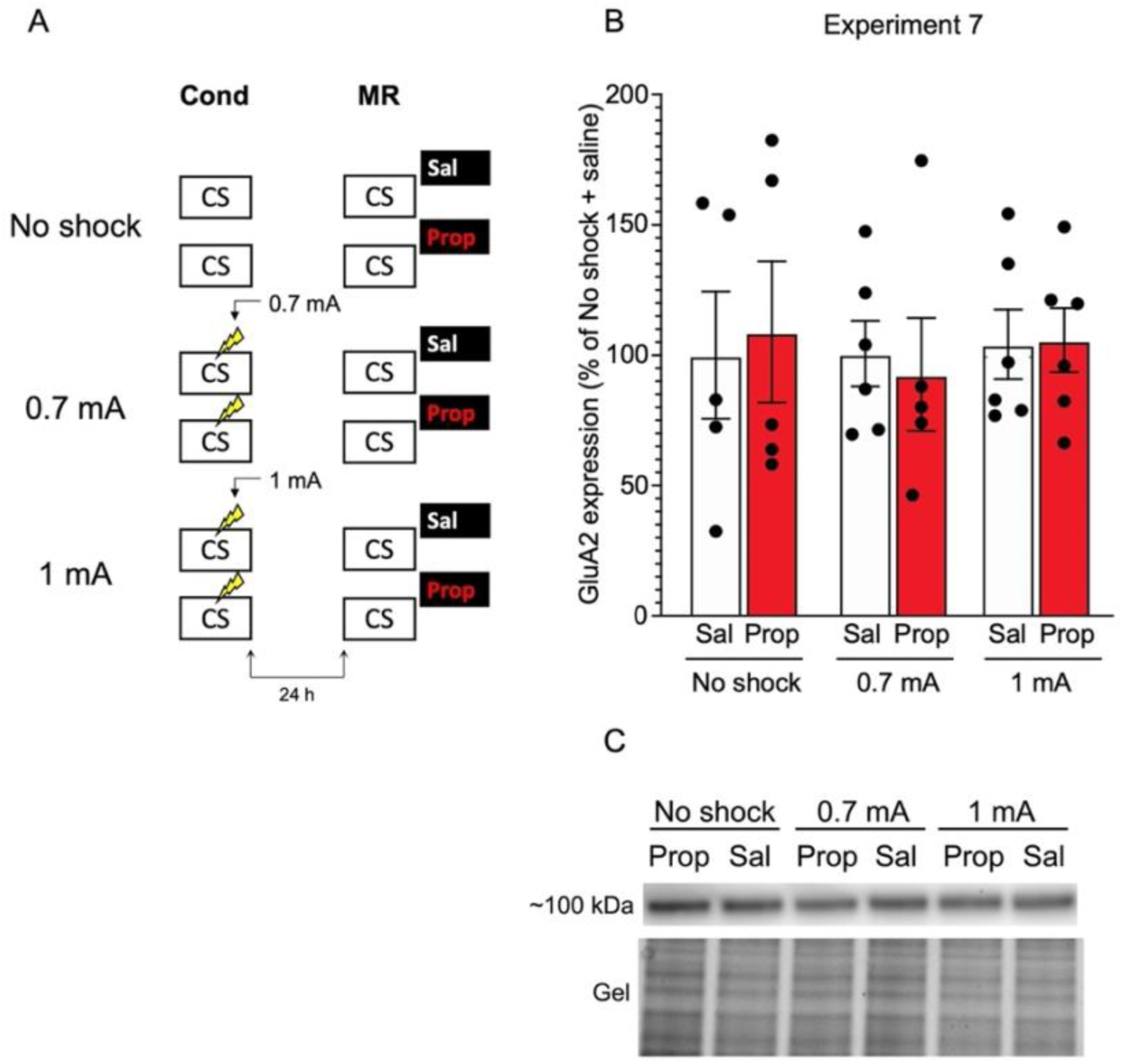
No effect of reactivation on synaptic GluA2 expression. (**A**) Design of Experiment 7. Mice received either no shock, one 0.7 mA shock, or one 1 mA shock during fear conditioning. One day later, mice underwent memory reactivation and received either saline (no shock, *n* = 5; 0.7 mA, *n* = 6; 1 mA, *n* = 6) or propranolol (no shock, *n* = 5; 0.7 mA, *n* = 5; 1 mA, *n* = 6) afterwards. One hour later the mice were sacrificed. Cond = conditioning, MR = memory reactivation, CS = conditioned stimulus (context), Sal = saline, Prop = propranolol. (**B**) Results of Experiment 7. GluA2 expression levels relative to the no shock + saline control group (saline in white bars, propranolol in red bars). Error bars represent SEM. Filled circles indicate individual animals. (**C**) Upper panel: Representative immunoblots for GluA2 with approximate molecular weight indicated. Lower panel: corresponding gels of total protein that were used for normalization.

### 3.7. Experiment 8

To successfully trigger memory destabilization, it is imperative to have robust memory acquisition and reactivation. In most of the experiments conducted so far, freezing levels during memory reactivation varied greatly, with maximum freezing levels around 60–80% and minimum freezing below 10% within single experiments. Hence, mice may have differred substantially in the extent to which their fear memories were reactivated and destabilized, such that reconsolidation was triggered in only a subset of the subjects, leading to small effects at best. In the next experiments, we therefore aimed to test training protocols that would lead to relatively homogenous freezing during reactivation across subjects. As learning by repetition is one of the most classic and widely-known principles of memory (Ebbinghaus, 2013), we administered multiple USs at regular intervals, instead of a single foot shock, as a means to induce robust fear learning in all animals. In Experiment 8, we presented three 0.5 mA shocks during conditioning, a procedure that has been shown to result in a fear memory that is vulnerable to post-reactivation administration of amnestic agents (Schmidt et al., 2017) (Figure 8A). We included both propranolol and anisomycin conditions. Again, no amnestic effect of propranolol or anisomycin was observed (Drug × Session, *F*2,24 = 0.205, *p* = 0.816, Figure 8B).

**Figure. 8.**
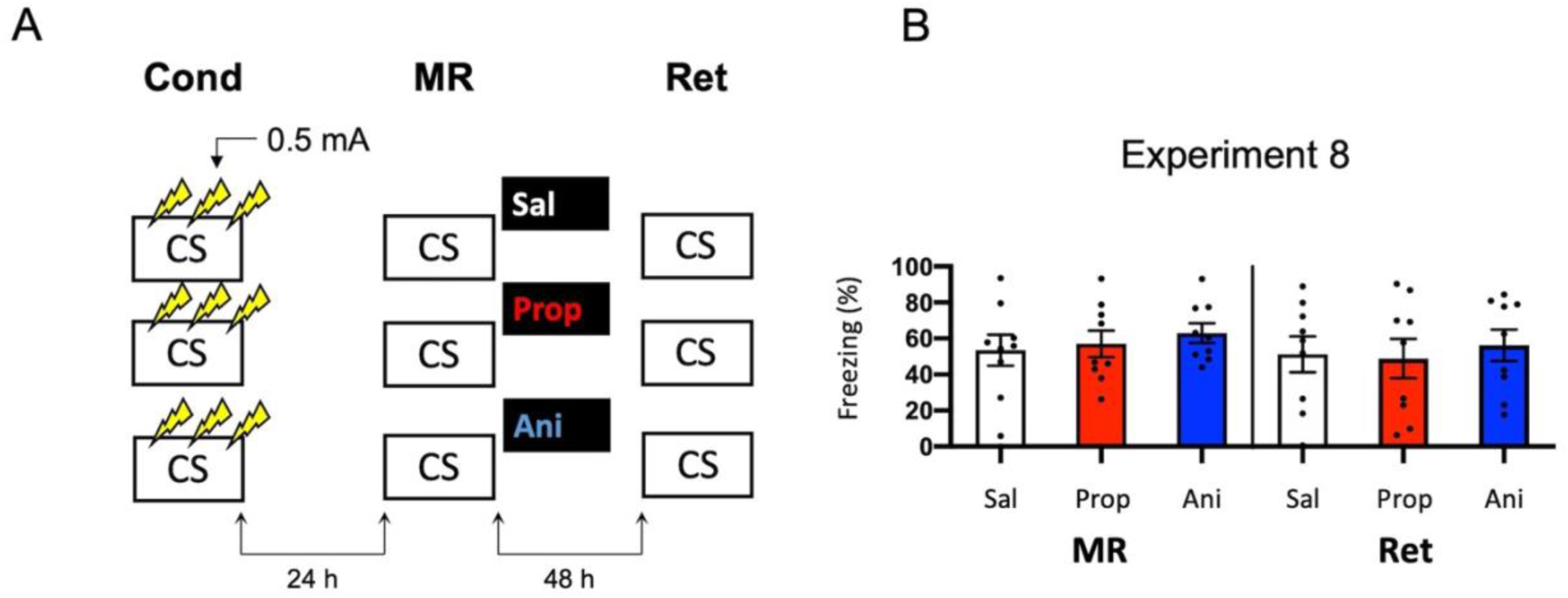
No effect of propranolol or anisomycin on reactivated memories that were acquired with three 0.5 mA shocks during contextual fear conditioning. (**A**) Design of Experiment 8. Mice received three 0.5 mA shocks during fear conditioning. One day later, mice received saline (*n* = 9), propranolol (*n* = 9), or anisomycin (*n* = 9) after memory reactivation. Forty-eight hours afterwards, retention of contextual fear was assessed. Cond = conditioning, MR = memory reactivation, Ret = retention, CS = conditioned stimulus (context), Sal = saline, Prop = propranolol, Ani = anisomycin. (**B**) Results of Experiment 8. Average percentage of freezing during memory reactivation is displayed on the left, and freezing during the retention test on the right panel of the column chart (saline in white bars, propranolol in red bars, and anisomycin in blue bars). Error bars represent SEM. Filled circles indicate individual animals.

### 3.8. Experiment 9

Note that the freezing levels during memory reactivation were significantly higher in Experiment 8, which involved three 0.5 mA shocks, than Experiment 6, in which a single 1 mA shock was administered (*t*50 = -7.161, *p* < 0.001, compare Figures 6B and 8B). Previous research has shown that increasing the number of administered shocks in contextual fear conditioning leads to higher freezing during a subsequent retention test (Poulos et al., 2016). Therefore, it is possible that multiple USs of a lower intensity results in a stronger fear memory than a single higher intensity US. The freezing levels during memory reactivation in Experiment 8 (three 0.5 mA shocks), relative to Experiment 6 (one 1 mA shock), were enhanced to such a striking extent (Cohen’s *d* = 1.99) that memory strength may have prevented the occurrence of diminished fear responding by administration of amnestic agents upon memory reactivation (Suzuki et al., 2004). In a final attempt to observe post-reactivation amnesia, we aimed to obtain relatively homogenous freezing using multiple shocks, but at the same time not induce a fear memory that is much stronger than in the previous experiments that included a single shock. Specifically, we repeated Experiment 8 and lowered the shock intensity to 0.3 mA (Experiment 9a) (Figure 9A).

**Figure. 9.**
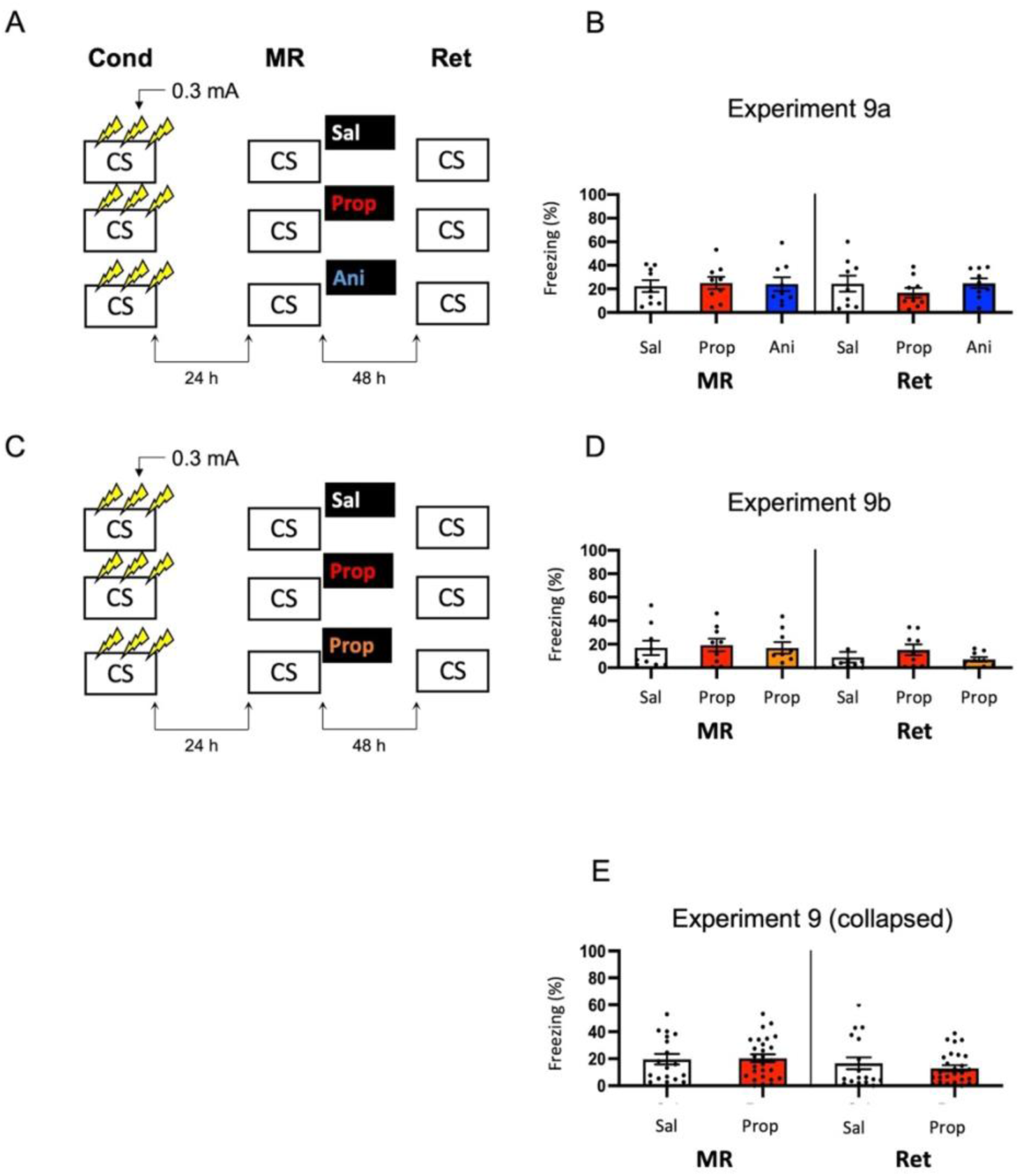
No effect of propranolol or anisomycin on reactivated memories that were acquired with three 0.3 mA shocks during contextual fear conditioning. (**A**) Design of Experiment 9a. Mice received three 0.3 mA shocks during fear conditioning. One day later, mice received saline (*n* = 9), propranolol (*n* = 9), or anisomycin (*n* = 9) after memory reactivation. Forty-eight hours afterwards, retention of contextual fear was assessed. Cond = conditioning, MR = memory reactivation, Ret = retention, CS = conditioned stimulus (context), Sal = saline, Prop = propranolol, Ani = anisomycin. (**B**) Results of Experiment 9a. Average percentage of freezing during memory reactivation is displayed on the left, and freezing during the retention test on the right panel of the column chart (saline in white bars, propranolol in red bars, and anisomycin in blue bars). (**C**) Design of Experiment 9b. The anisomycin condition was replaced with propranolol dissolved in saline at half the usual ratio and injected in double the volume/body weight (*n* = 9 in all conditions). All other procedures were the same as Experiment 9a. (**D**) Results of Experiment 9b (saline in white bars, propranolol in red bars, and propranolol double volume in orange bars). (**E**) Collapsed results of Experiment 9a and 9b (propranolol in all volumes). Error bars represent SEM. Filled circles indicate individual animals.

Once again, no evidence for a post-reactivation amnestic effect of propranolol or anisomycin was found (Drug × Session, *F*2,24 = 1.227, *p* = 0.311, Figure 9B). When animals that showed freezing levels below 10% during reactivation were excluded, a Drug x Session interaction at trend level was found (*F*2,17 = 3.214, *p* = 0.065, supplementary Figure S1L). Additional independent t-tests showed that freezing during the retention test was lowered in the propranolol condition, relative to the saline condition (*t*11 = 2.264, *p* = 0.045). Surprisingly, this was not the case in the anisomycin condition (*t*11 = 0.645, *p* = 0.532). Although a reduction in fear responding by post-reactivation administration of propranolol is in line with earlier reports on the involvement of β-ARs in memory reconsolidation (Johansen et al., 2011), the results of Experiment 9a are hard to reconcile with the presumed protein dependency of this process (Nader et al., 2000). Integration theory as an alternative account, also does not provide an explanation, as there is no indication that propranolol would, but anisomycin would not, induce a state-dependent memory (Gisquet-Verrier and Riccio, 2018). Furthermore, as this reducing effect of propranolol on freezing levels from memory reactivation to retention test was only observed with the specific inclusion criterion of at least 10% freezing during reactivation, it is uncertain to what extent this finding is robust. We therefore performed a replication of the experiment (Experiment 9b) in which we aimed to maximize the chance of observing post-reactivation amnesia by administration of propranolol. Specifically, we included a condition in which propranolol was injected in the same concentration as in the previous experiment (2 mg propranolol/1 mL saline), and a condition in which half the concentration and double the volume was injected like in Experiment 3 (Villain et al., 2016)) (Figure 9C). Thereby, two different administrations were used that might result in diminished fear responding.

No reduced fear responding by administration of propranolol was observed when all mice were included (Drug × Session, *F*2,24 = 0.737, *p* = 0.489, Figure 9D), nor when mice that showed freezing levels below 10% were excluded (Drug × Session, *F*2,13 = 1.632, *p* = 0.233, Figure S1M). Since in experiment 9b a large number of mice showed freezing levels below 10%, there was low statistical power to replicate the propranolol-induced reduction in fear responding that was observed in Experiment 9a (1-β = 0.40). We, therefore, also analyzed collapsed data of Experiment 9a and 9b. Both with (*F*1,43 = 1.261, *p* = 0.268, Figure 9E) and without (*F*1,27 = 2.249, *p* = 0.145, Figure S1N) inclusion of mice that showed freezing levels lower than 10% during reactivation, no effect of propranolol (in any volume) was found, suggesting that the post-reactivation amnesia as observed in Experiment 9a was not robust. Furthermore, a cross-experimental comparison (Experiment 9 *vs*. Experiment 6, compare Figures 9E and 6B) showed that administering multiple shocks, instead of a single shock, did not lead to more homogenous freezing during reactivation (Levene’s test, *F*1,68 = 1.313, *p* = 0.256), which was the aim of the last two experiments. Thus, as the different behavioral protocols involving single or multiple foot shocks of varying intensities did not provide a stable paradigm to study post-reactivation amnesia, and no clear indications of how to improve the protocol seemed to be at hand, no further experiments were carried out.

## 4. Discussion

We performed a series of experiments to study the effects of post-reactivation, systemic administration of propranolol on contextual fear memory in mice. The eventual purpose of this study was to elucidate the neural mechanisms by which this β-AR antagonist has been shown to interfere with contextual fear memories (Abrari et al., 2008; Liu et al., 2015), and test ways to overcome boundary conditions that potentially complicate application of post-reactivation amnesia in clinical practice (Kindt, 2014). However, in a total of 12 experiments (including replications) that involved different fear acquisition procedures, drugs (i.e., not only propranolol, but also anisomycin), and read-outs of post-reactivation amnesia (i.e., behavioral, cellular, molecular), our attempts to observe robust amnesic effects were largely unsuccessful. At best, only slight reductions in freezing were found in a minority of the experiments.

Specifically, only when the data of Experiments 2a and 2b were collapsed, a marginally significant effect of post-reactivation administration of propranolol on freezing was observed. Note that in these experiments we attempted to create stronger fear memories relative to Experiment 1 by raising the intensity of the delivered foot shock. It is remarkable that we indeed found an indication of reduced fear responding in the propranolol condition in Experiment 2, even though the freezing levels during memory reactivation were not significantly higher in this experiment than in Experiment 1. As it may seem unlikely that experiments with comparable levels of freezing during memory reactivation sometimes do (Experiments 2), and sometimes do not (Experiment 1) show pharmacologically-induced amnesia after memory reactivation, one could question whether the finding in Experiment 2 is a true positive. Previous research has, however, indicated that the expression of fear during memory reactivation is not an accurate index of memories entering a labile state. Emotional memory expression during memory reactivation is not a necessary, nor a sufficient condition for the occurrence of post-reactivation amnesia (see Faliagkas et al., 2018 for a review). Despite the freezing levels in Experiments 1 and 2 being similar, the memory processes that were triggered by reactivation were therefore not necessarily identical. Nevertheless, the effect in Experiments 2 was considerably smaller than in earlier demonstrations of reduced fear responding by administration of propranolol after memory reactivation (e.g., Przybyslawski et al., 1999; Dȩbiec and Ledoux, 2004; Liu et al., 2015), and the small effect did not replicate in two additional experiments (Experiments 3 and 6). The minor amnestic effect also did not translate to a difference in neural activity between experimental groups. All in all, even though the results of Experiments 2 were perhaps not a chance finding, the observed reduction in fear responding by post-reactivation administration of propranolol was clearly not robust either.

How can we explain that our attempts to develop a paradigm that would show consistent amnestic effects were unsuccessful, given that we tried a wide variety of procedures? A few factors could be considered. The null findings do not seem to be attributable to a lack of power. The number of mice that were included in each experiment were comparable to previous studies in which large effects were observed (e.g., Przybyslawski et al., 1999; Dȩbiec and Ledoux, 2004; Liu et al., 2015). We also did not find robust effects when the data of similar experiments (Experiment 1 and 2, Experiment 3 and 4, Experiment 3 and 6) or all experiments performed within the same research instrument (TSE, or Med Associates, see supplementary results) were collapsed. Furthermore, a general inefficiency of propranolol in interfering with fear memories does not seem to be a plausible explanation either. We administered propranolol in the same dose as in previous experiments that have shown robust post-reactivation amnesia in both rats (Przybyslawski et al., 1999; Dȩbiec and Ledoux, 2004) and mice (Liu et al., 2015; Villain et al., 2016), and in two different volumes of saline, of which one has been used earlier in mice (Villain et al., 2016). We also included anisomycin in several experiments, which in previous research has induced amnestic effects across various protocols. For example, reduced fear responding by local or systemic administrations of anisomycin before or after reactivation of contextual or cued fear have all been observed (e.g., Nader et al., 2000; Lee et al., 2004; Suzuki et al., 2004; Parsons et al., 2006; Blundell et al., 2008; Rao-Ruiz et al., 2011; Kwak et al., 2012). Finally, we found no effects of memory reactivation on expression of the glutamate receptor subunit GluA2 (i.e., a marker of memory destabilization, Rao-Ruiz et al., 2011; Bhattacharya et al., 2017). This is in line with recent research showing similar null findings in rats using cued fear conditioning and Shank protein as a marker of memory destabilization (Rotondo et al., 2022, see also Luyten et al. (2021) for more behavioral data).

The most plausible explanation for the null findings is, therefore, that the procedures did not (consistently) trigger the memory to enter a state in which it becomes receptive to change. Previous research has identified several boundary conditions for reactivation-dependent memory plasticity (Pedreira et al., 2004; Lee, 2009). For example, both memories that are reactivated rather shortly or lengthily (Sevenster et al., 2012, 2014; Merlo et al., 2014), in relation to their strength (Suzuki et al., 2004), may not undergo weakening by administration of amnestic agents. At first sight, it seems like we have largely ruled out this explanation, since we have varied the fear-conditioning procedures across experiments to induce memories of different strengths. However, the experiments showed highly variable levels of freezing during memory reactivation, ranging from lower than 5% to higher than 85% within a single condition in the most extreme cases. It therefore remains possible that memories became vulnerable to the administration of propranolol or anisomycin in only a small number of mice (although in some previous studies post-reactivation amnesia was observed with somewhat similar spread of freezing during reactivation, Dȩbiec et al., 2002; Rao-Ruiz et al., 2011). Since boundary conditions can complicate falsifiability of any phenomenon (Elsey and Kindt, 2017), it is important to test such explanations before final conclusions are drawn. For this reason, we have performed experiments (Experiment 8–9) that were specifically aimed at reducing the spread of freezing during memory reactivation. These experiments were however unsuccessful in this regard. Whether inconsistency of memory destabilization is the cause of the present (or other) null findings thus remains an open question at this point. Future studies aimed at pharmacologically potentiating memory destabilization, for instance by using partial NMDA receptor agonists, could help clarify this point (Bustos et al., 2010).

Another possibility to consider, although speculative, is that the inbred experimental mouse strain used by us has undergone changes over the past years, such that procedures that used to be effective no longer trigger a state of memory plasticity (Schroyens et al., 2019). It is remarkable in this light that Experiment 1 was procedurally identical to an earlier study (Rao-Ruiz et al., 2011) in many respects (animal strain, supplier, lab space, behavioral system, training protocol and length of memory reactivation and retention tests), but now showed much lower freezing levels during reactivation. This discrepancy might be explained by previous research showing that inbred mouse strains are not always isogenic. Factors such as genetic drift, spontaneous mutations and epigenetic changes may all influence the behavior of experimental animals (Stiedl et al., 1999; Loos et al., 2015; Oey et al., 2015; Chebib et al., 2021). We tried to control for this variability by ordering animals from a well-established breeder (Charles River laboratories, France) that has robust genetic monitoring programs in place. However, although the commonly used genetic screens of commercial breeders minimize cross-strain contamination, they are less successful at identifying spontaneous mutations and epigenetic changes that may drive behavioral variability without presenting as overt phenotypes. Isogenicity within the inbred strain can therefore not be assumed (Chebib et al., 2021).

Besides genetic variability, environmental factors may also contribute to the limited replicability of reactivation-induced amnesia that we observe. Factors such as breeding conditions, colony maintenance, maternal care, or transportation, which are all beyond the researchers’ sight or control, might impact the success of downstream experimental manipulations. For example, early life stress, which could potentially occur or vary among experimental animals before their arrival at the research site, has been shown to have an influence on both fear learning (Kosten et al., 2012) and memory plasticity (Villain et al., 2018; Couto-Pereira et al., 2019). Variations in maternal care and colony maintenance can also affect adult behavioral outcomes and stress reactivity of C57Bl6/J mice (Pedersen et al., 2011).

When trying to explain the varying freezing levels across studies, a possibility to consider is that genetic factors, environmental factors, or an interaction thereof, which have been shown to contribute to within-strain variability (Loos et al., 2015), result in a change in risk assessment of aversive stimuli and a choice in downstream defensive strategies by experimental animals. Aversive learning can contain explicit and implicit memory systems that are not readily dissociable in rodents (Wotjak, 2019). This is attributable to a lack of adequate readouts that can distinguish between behavioral outcomes driven by associative and non-associative components of the fear circuit (Wotjak, 2019). Automated fear-conditioning systems, like the ones used in this study, measure freezing as the sole behavioral output of fear. However, in addition to vegetative (e.g. changes in heart rate, respiration) and hormonal (e.g. changes in stress hormone levels) responses, rodents can use risk assessment to choose an appropriate behavioral defensive response to threatening stimuli that they encounter (Blanchard et al., 2011). This analytic function enables animals to choose a strategy most likely to succeed in a given situation and include freezing, flight/escape, startle, burying, ultrasonic vocalizations, or a combination thereof (Wotjak, 2019). A switch in defensive strategy, such as active ‘flight’ responding instead of a cessation of movement, would result in generally lower freezing levels that could potentially affect behavioral readouts and confound the interpretation of the amnestic effects of the applied interventions. The particularly low freezing in Experiment 4 in which the experimental procedures were executed in the animals’ dark phase suggests that mice can indeed use different of such defensive strategies in response to the same aversive stimuli. Similar to our arguments concerning high variability in memory reactivation, the influence of genetic and environmental factors might prove to be difficult to control, and in that case remain unfalsifiable. However, being able to breed and maintain inbred colonies in-house or change to paradigms that call upon more active defensive responses, such as inhibitory avoidance, could shed light on some of these ideas.

In the present series of experiments, no direct replication attempts of earlier studies have been performed, which has been done in some experiments of Schroyens et al. (2019.) The findings presented here do not suggest that previously reported results are false positives. Due to drastic changes in the animals’ behavior as discussed above, the overarching approach towards a paradigm with consistent effects was to gradually build on the outcomes of individual experiments in the series. Although we have not been successful in doing so, we believe that this might be the most fruitful way forward. Apart from replication studies showing that seemingly basic manipulations do not always have consistent outcomes, systematically varying single factors such as the strength and/or the number of the aversive stimulus presentations, as well the duration and time-point of reactivation, could contribute to regaining control over reactivation-dependent manipulation of memory in a step-by-step fashion. Our findings add to this endeavor, specifically in relation to contextual fear memories in mice. Apart from freezing, we did not observe effects of post-reactivation administration of propranolol at a neural level either, which is reminiscent of a recent study by Leal Santos et al. (2021). These authors demonstrated that propranolol only has an acute effect on contextual memory expression and DG memory trace cells when delivered immediately prior to memory reactivation. No long-term effect on memory retention after drug washout (i.e., 24 h later) was observed. The authors –like us – observed no effect of post-reactivation administration of propranolol on a subsequent retention test, while using a different mouse strain (129S6/SvEv) and conditioning protocol (4 × 0.75 mA foot shocks) than in the present study. Together, our data and those of Leal Santos et al. (2021) thus seem to suggest that blockade of β-ARs does not have lasting effects on neural activity in the hippocampal DG.

In conclusion, our findings show that inducing reactivation-dependent reductions in contextual fear responding is not as straightforward as the overall state of the published literature on reconsolidation suggests, yet align with a recent report of null findings in rats (Schroyens et al., 2019). As we used a different species, laboratory equipment, behavioral protocols, and drugs, incorporated read-outs of memory plasticity at several levels (behavioral, cellular, molecular), and included larger samples in each experiment than in Schroyens et al., 2019, the findings show that obstacles in observing post-reactivation amnesia are not uniquely related to the parameters varied by particular research groups. The present findings do not rule out the existence of post-reactivation amnesia. They do underscore the elusive nature of this phenomenon and emphasize the need for better control over and understanding of the intrinsic and extrinsic factors that may govern it.

## Author contributions

**Wouter Cox:** Conceptualization, Formal Analysis, Investigation, Data Curation, Writing- Original Draft Preparation, Visualization. **Leonidas Faliagkas:** Conceptualization, Formal Analysis, Investigation, Data Curation. **Amber Besseling:** Investigation. **Rolinka van der Loo:** Investigation. **Sabine Spijker:** Conceptualization, Writing- Reviewing and Editing, Funding Acquisition. **Merel Kindt:** Conceptualization, Writing- Reviewing and Editing. **Priyanka Rao-Ruiz**: Conceptualization, Methodology, Formal Analysis, Investigation, Resources, Writing- Reviewing and Editing, Visualization, Supervision, Project Administration, Funding Acquisition.

