## Supplementary data for "Interfering with contextual fear memories by post-reactivation administration of propranolol in mice: a series of null findings"

### *Supplementary Material*

### 1. Supplementary Results

To rule out that the null findings in the present study could be explained by insufficient statistical power within the individual experiments, we performed additional analyses with the data of multiple experiments collapsed. All experiments were sorted by research instrument (TSE systems, Med Associates system), and drug (saline, propranolol, anisomycin). As Experiment 5 involved a different reactivation duration than all other experiments and Experiment 7 did not involve a retention test, these two experiments were excluded from this analysis. Across the experiments performed in a TSE system (Experiments 1–2), there was no effect of post-reactivation administration of propranolol versus saline on changes in freezing from memory reactivation to retention test (Drug × Session, $F_{1,70} = 0.187$, $p = 0.667$). For the collapsed data of experiments that were performed in a Med Associates system (Experiment 3–4, 6, 8, 9), this same analysis also showed no significant effect ($F_{1,129} = 1.732$, $p = 0.190$). Finally, when the data of all anisomycin conditions were collapsed (Experiment 6, 8, 9), no effect was found either ($F_{1,50}$ = 0.394, $p$ = 0.533). Therefore, low statistical power does not seem to be an explanation of the null findings in the present series of experiments.

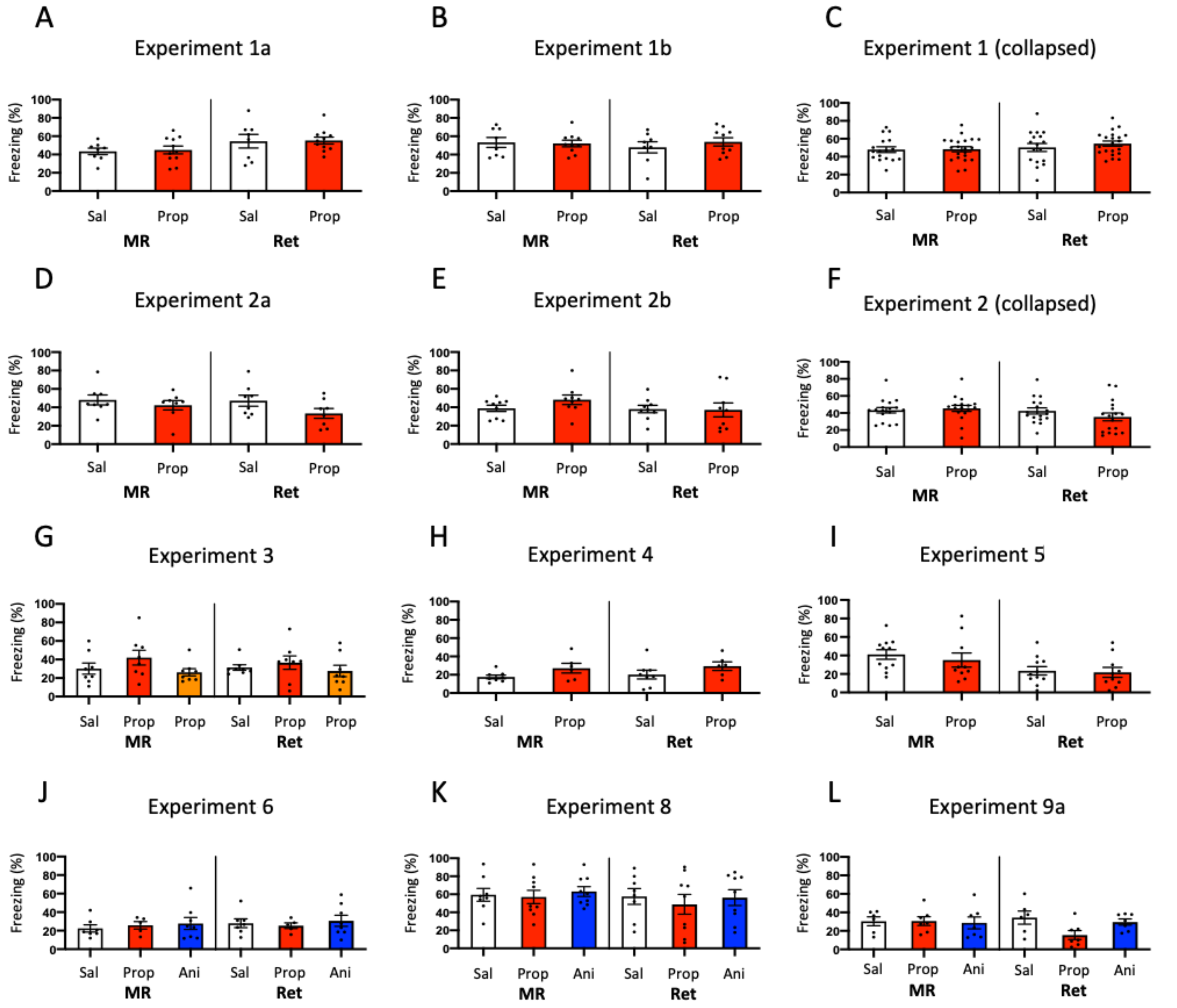


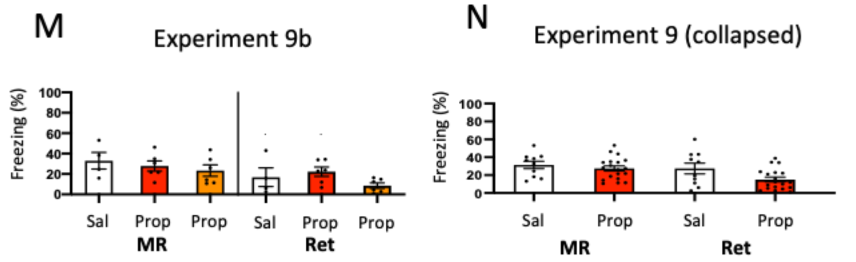


**Fig. S1.** Analyses of Experiment 1–6, 8, and 9 in which animals that showed lower than 10% freezing during memory reactivation were excluded. Cond = conditioning, MR = memory reactivation, Ret = retention, Sal = saline, Prop = propranolol, Ani = anisomycin. (**A**) Experiment 1a. One mouse was excluded due to low freezing (propranolol condition). No Drug x Session interaction was found, $F_{1,17}$ = 0.008, $p$ = 0.929. (**B**) Experiment 1b. No mice showed freezing levels lower than 10% during memory reactivation. (**C**) Collapsed data of Experiment 1a and 1b. No Drug x Session interaction was found, $F_{1,35} = 0.270$, $p = 0.607$. (**D**) Experiment 2a. No mice showed freezing levels lower than 10% during memory reactivation. (**E**) Experiment 2b. No mice showed freezing levels lower than 10% during memory reactivation. (**F**) Collapsed data of Experiment 2a and 2b. (**G**) Experiment 3. Three mice were excluded (saline condition, *n* =1; propranolol condition, *n* = 1, propranolol double volume condition, *n* = 1). No Drug x Session interaction was found, $F_{2,21} = 0.437$, $p = 0.652$. (**H**) Experiment 4. Ten mice were excluded due to low freezing (saline condition, *n* = 4; propranolol condition, *n* = 6). No Drug x Session interaction was found, $F_{1,12} = 0.002$, $p = 0.963$. (**I**) Experiment 5. One mouse was excluded (propranolol condition). No Drug x Session interaction was found, $F_{1,19} = 0.036$, $p = 0.852$. (**J**) Experiment 6. Five mice were excluded (saline condition, *n* =1; propranolol condition, *n* = 4). No Drug x Session interaction was found, $F_{2,17} = 0.422$, $p = 0.662$. (**K**) Experiment 8. One mouse was excluded (saline condition). No Drug x Session interaction was found, $F_{2,23} = 0.207$, $p = 0.815$. (**L**) Experiment 9a. Seven mice were excluded (saline condition, *n* = 3; propranolol condition, *n* = 2; anisomycin condition, *n* = 2). A Drug x Session interaction was observed at trend level, $F_{2,17} = 3.214$, $p = 0.065$. (**M**) Experiment 9b. Eleven mice were excluded (saline condition, *n* = 5; propranolol condition, *n* = 3; propranolol double volume, *n* = 3). No Drug x Session interaction was found, $F_{2,13} = 1.632$, $p = 0.233$. (**N**) Collapsed data of Experiment 9a and 9b. No Drug x Session interaction was found, $F_{1,21} = 1.151$, $p = 0.295$. Error bars represent SEM. Filled circles indicate individual animals.

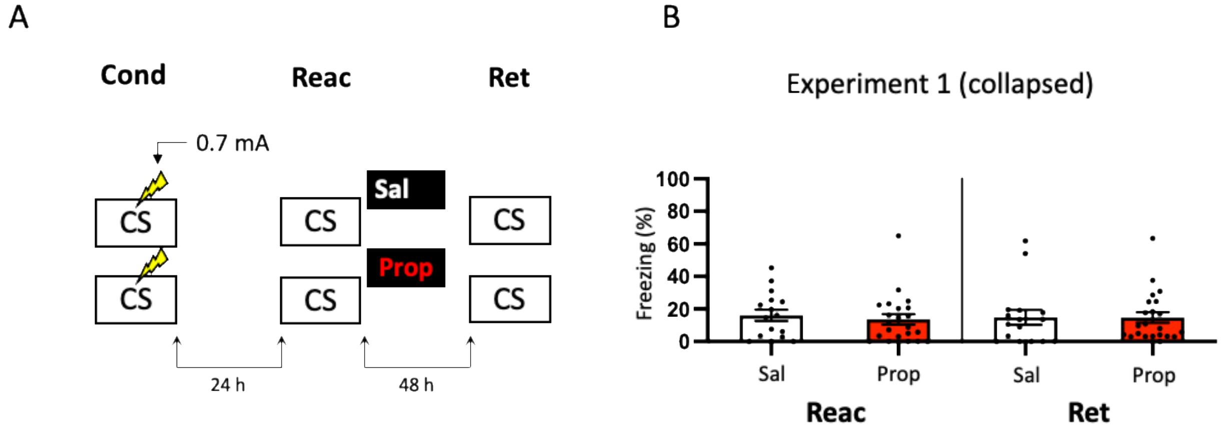


**Fig. S2**. Results of Experiment 1 with freezing measured using a time threshold of 5 s, as in a previous study that used the same protocol (Rao-Ruiz et al., 2011). Cond = conditioning, MR = memory reactivation, Ret = retention, CS = conditioned stimulus (context), Sal = saline, Prop = propranolol. Error bars represent SEM. Filled circles indicate individual animals. Again, no effect of post-reactivation administration of propranolol was found on contextual fear ($F_{1,36}$ = 0.126, $p$ = 0.725). Most importantly, freezing levels during memory reactivation are notably lower than in this previous study (Rao-Ruiz et al., 2011, Fig. S1D).
